# High quality mapping of chromatin at or near the nuclear lamina for small numbers of cells

**DOI:** 10.1101/2021.01.03.425156

**Authors:** Joseph R. Tran, Xiaobin Zheng, Stephen A. Adam, Robert D. Goldman, Yixian Zheng

## Abstract

The chromatin associated with the nuclear lamina (NL) is referred to as Lamina-Associated Domains (LADs). While mapping of this feature has been done using various technologies, technical limitations exist for each of the methods. Here, we present an adaptation of the Tyramide-Signal Amplification sequencing (TSA-seq) protocol, which we call chromatin pull down-based TSA-seq (cTSA-seq), that can be used to map chromatin regions at or near the NL from as little as 50,000 cells without using carriers. The cTSA-seq mapped regions are composed of LADs and smaller chromatin regions that fall within the chromatin B-compartment known to be enriched for heterochromatin and be present at the nuclear periphery. As a proof of principle, we used cTSA-seq to map chromatin at or near the assembling NL as cells exit mitosis and progress through early and later G1. Consistent with previous reports, lamin-B1 based cTSA-seq revealed that regions toward the distal ends of chromosomes are near or at the reassembling NL during early G1. The cTSA-seq mapping and analyses revealed similarity between the early G1 chromatin and oncogene-induced senescent cell populations. The cTSA-seq reported here represents a useful method for analyzing chromatin at or near the NL from small numbers of cells.

## Introduction

A portion of the metazoan genome is organized into compact, transcriptionally repressed DNA known as heterochromatin. In most cell types, a major fraction of heterochromatin is localized to the nuclear periphery and may directly interact with a protein meshwork known as the nuclear lamina (NL) that contains the filamentous A- and B-type lamins (1). The heterochromatin that directly interacts with the NL is known as Lamina-Associated Domains (LADs) (2). A comparison of LADs and the A/B compartments defined from Hi-C experiments showed that LADs are in the B compartment, which is defined as “closed” chromatin (2-4). The A and B compartments are, however, complex and each compartment can be further divided into subcompartments. For example, the B compartment can be divided into sub-compartments B1 through B4 that are differentially enriched for epigenetic marks (e.g., H3K27me3 and H3K9me3) (4). Of these B subcompartments, the B3 compartment shows the strongest correlation with LADs followed by the B2 compartment, which is correlated with both LADs and pericentromeric heterochromatin (4). The B1 compartment appears to be facultative heterochromatin, as it shows a positive correlation with H3K27me3 and negative correlation with H3K36me3, while the B4 compartment represents a rare (0.4% of the genome) definition that is present only on chromosome 19 (4). Additional studies are needed to reveal if any chromatin in B compartments not mapped by LADs are near the NL. Mapping methods that identify chromatin at or near the NL would facilitate these investigations.

LADs are known to vary according to cell type (5,6) and recent work reveals that LADs, while largely stable, show dynamics during the cell cycle (7,8). The greatest dynamism has been reported during early G1 with an initial enrichment of NL-chromatin contacts occurring at the distal ends of chromosomes (7). These observations are consistent with previous studies of nuclear envelope reformation on decondensing post-mitotic chromosomes (9-12) and the reported increased spatial dynamics of genomic features, particularly those of telomeres, in early interphase (13-20). Collectively, these observations suggest a model where the distal ends of chromosomes transiently localize near the assembling NL in early G1 reminiscent of a Rabl-like configuration followed by re-organization and formation of a stable and typical LADs configuration.

LADs have been mapped by a variety of methods including chromatin-immunoprecipitation (ChIP)-seq, DNA adenine methyltransferase (Dam) identification (DamID) and Tyramide Signal Amplification (TSA)-sequencing (TSA-seq) (2,21-24). LADs were first mapped by DamID, which uses a fusion protein of bacterial Dam and lamin-B1 expressed at low levels in tissue culture cells (2). DNA located at the NL is methylated by contacting the Dam enzyme and can be identified by sequencing or microarray technologies. A modified form of this method can also be used to visualize LADs (19). The TSA-seq method employs a different strategy and uses antibody staining (e.g., anti-lamin-A/C) to specifically label the NL in fixed cells. A corresponding horseradish peroxidase (HRP)-conjugated secondary antibody then catalyzes the covalent attachment of biotin onto nearby proteins and DNA (21). The TSA-seq method uses a genomic DNA input where the fraction of biotinylated DNA is isolated by Streptavidin-bead pull down, purified and then subject to sequencing. Considering the diffusive nature of the TSA reaction, this method is useful both as a mapping tool and as a “cytological ruler” and variations of this method have also found utility in proteomics (21,25).

We recently adapted the use of the engineered Ascorbate Peroxide 2 (APEX2) enzyme as a genome mapping tool, which we termed APEX2-identification (APEX2-ID), to identify stable and variable chromatin regions at or near the NL during the cell cycle (8,26,27). The APEX2-ID method uses the highly active APEX2 enzyme to catalyze the deposition of biotin onto proteins near the NL-localized fusion protein, which can then be isolated after chromatin fragmentation by pull down using Streptavidin beads. APEX2-ID can also be used to map NL-proximal proteins and associated RNAs from live cells and the activity of this enzyme persists under different conditions including cellular fixation. The van Steensel group recently developed an approach called protein-A DamID (pA-DamID) to map LADs from different cell cycle stages (7). The pA-DamID approach leverages antibody targeting, in a manner similar to TSA-seq, of DamID and revealed LADs dynamics during the cell cycle. The approach also revealed the enrichment of NL-chromatin interactions toward the distal ends of chromosomes in the early stage of G1. The pA-DamID and TSA-seq approaches highlight the utility of antibody-based methods.

Pros and cons exist for these aforementioned methods. For instance, the APEX2-ID method is a multi-purpose tool useful for mapping genomic regions, proteins and RNAs and the APEX2 reaction can be completed in a matter of seconds, thus allowing high-temporal resolution. The APEX2-ID approach is, however, limited by transfection issues and unwelcomed overexpression effects. DamID is a classic straightforward method that directly analyzes DNA, and is the only LADs mapping method for which single cell mapping has been reported (19,28). However, DamID requires transfection/transduction and several hours to sufficiently methylate DNA as the fusion protein is expressed at low levels (28). The TSA-seq approach, as described by the Belmont group, is attractive as it allows the user to temporally resolve events based on the timing of cellular fixation and employs common immunostaining approaches familiar to most labs (21). The method, however, uses genomic DNA as an input source, and the fraction of DNA that is biotinylated was reported to be low (21), Thus, TSA-seq requires a large number of cells (>10 million) to obtain sufficient material and is not easily implemented in studies with small numbers of cells. Ultimately, a method that can reliably identify various genomic structures from a lower number of cells is needed as it opens the possibility of mapping from *in vivo* sources where cell numbers are low.

Of these existing methods, TSA-based labeling has a number of other distinct advantages. For example, it allows labeling of essentially any endogenously expressed protein of interest as has been shown for the mapping of both LADs and nuclear speckles (21). The TSA reaction is time efficient, available as a commercial kit used regularly for tissue section staining, and it enables the downstream processing that takes advantage of the simplicity and familiarity of the pull down approach commonly used in ChIP-seq experiments.

Based on our experience using the APEX2-ID method to identify chromatin regions at or near the NL by Streptavidin-based pull down of chromatin, we explored the use of a chromatin pull down-based version of TSA-seq, which we call cTSA-seq. The cTSA-seq method reported here can map chromatin regions at or near the NL from 50,000 cells without the addition of carrier. It can sensitively identify both LADs and additional NL-proximal regions that are heterochromatic and found in the B compartments. As a proof of principle, the cTSA-seq approach can be used to identify chromatin interactions at or near the reassembling NL during early G1 as previously reported (7). We further note that the early G1 chromatin-NL relationship is similar to those mapped from the oncogene-induced senescence cells (29,30).

## Materials and Methods

### Reagents

The Biotin-XX-Tyramide Superboost kit (Invitrogen, USA, #2005939) was used for the TSA reaction. The rabbit anti lamin-B1 antibody (Abcam, USA, #ab16048) and a Streptavidin conjugate to Alexa 488 (1:200, Biolegend, #405235) were used for fluorescence staining.

### Biological resources and culturing conditions

HCT116 (ATCC, #CCL-247), K562 (ATCC, #CCL-243), hTERT-RPE-1 (ATCC #CCL-4000) and mouse embryonic fibroblasts (MEFs) were cultured in McCoy’s 5a with 10% FBS (HCT116), IMDM media with 15% (K562), DMEM-F12 with 15% (hTERT-RPE-1), and DMEM supplemented with 15% FBS, respectively. HCT116, hTERT-RPE-1 and MEFs were cultured in 100 or 150mm dishes. K562 cells were cultured in T25 flasks. All cells were cultured at 37 °C in 5% CO_2_. Bulk cell populations were counted using a hemacytometer and experiments with lower cell numbers were performed by serially diluting of the harvested cells.

### Nocodazole block, release and harvest for early and later G1 cells

HCT116 or hTERT-RPE-1 cells were cultured in three 150mm dishes to ∼40-60% confluency per experiment and then exposed to nocodazole (100ng/ml) for 18 hours. The cells were washed two times with 10 ml of pre-warmed media and mitotic cells were shaken off and seeded onto 100mm dishes that were pre-coated with poly-lysine. A fraction of the shake-off cells was saved and fixed with 1% paraformaldehyde for 10 minutes at room temperature to check for purity of the shake-off population by FACS (see below). The seeded cells were harvested at the indicated post-seeding times by trypsinization and then immediately fixed with 1% paraformaldehyde for 10 minutes at room temperature. For experiments with G1, S, and G2 phases, asynchronous cultures were harvested by trypsinization and fixed for 10 minutes with 1% paraformaldehyde. Paraformaldehyde fixation was neutralized with 125mM glycine and the cells were pelleted at 300g x 5 minutes and washed with phosphate buffered saline (PBS).

### Fluorescence Activated Cell Sorting (FACS) to analyze and isolate cells at different stages of cell cycle

All FACS was performed on a BD FACSaria III machine and on average we obtained 800,000 to 1 million cells unless noted otherwise. To isolate asynchronous cell cycle stage-specific populations (e.g., G1, S and G2), and the early or later G1 populations obtained after nocodazole block, cells were fixed as described above and the resuspended in HBSS + 2% FBS + 10μg/ml Hoechst 33342. The cells were incubated with Hoechst 33342 for at least 20 minutes at room temperature before FACS. For nocodazole block and release, we rejected samples where the shake-off mitotic population contained greater than 1% of contaminating G1 cells.

For SLAM-seq experiments (see section, “Thiol (SH)-Linked Alkylation for the Metabolic sequencing of RNA” below), FACS based cell isolation was done with live cells stained with 10μg/ml Hoechst 33342 in the last 20 minutes of culture. Since FACS was done with live cells, we limited sorting time to a maximum of 10 minutes in order to reduce the time between 4sU labeling and RNA harvest. Sorting for SLAM-seq experiments yielded between 30,000 to 100,000 cells.

### Chromatin pull down-based TSA-seq (cTSA-seq)

Paraformaldehyde (1%)-fixed cell populations were permeabilized with PBS containing 0.25% Triton X-100 for 10 minutes at room temperature. Permeabilized cells were pelleted at 200g for 5 minutes and the supernatant was carefully removed using a P100 pipette. The cells were then resuspended in PBS + 1% hydrogen peroxide and incubated for 20 minutes at room temperature to neutralize endogenous peroxidases. The suspension was then pelleted at 200g for 5 minutes and carefully aspirated with a P100 pipette. The cells were blocked for 20 minutes in PBS containing 10% normal goat serum, 10% BSA and 10mM sodium azide and pelleted as described above. The cells were resuspended by gently flicking the tube in the same blocking buffer containing 1:250 dilution of rabbit anti lamin-B1 antibody (Abcam, #ab16048) and incubated overnight at 4 °C in a microfuge tube rack. The following day, the cells were pelleted at 200g for 5 minutes and aspirated with a pipette. The pellet was washed 2 times for 15 minutes each with PBS supplemented with 0.001% Tween-20. The secondary antibody staining and TSA reaction reagents were from a commercial Biotin-XX-Tyramide Superboost kit (Invitrogen, #2005939) and were used as follows. The primary antibody stained cells were resuspended by gently flicking in ∼100μl of the provided anti-rabbit horseradish peroxidase (HRP) secondary antibody solution and incubated in a microfuge tube rack at room temperature for 2 hours with intermittent mixing. The cells were pelleted at 200g for 5 minutes, washed 3 times for 15 minutes each with PBS + 0.001% Tween-20 and one time with 1x reaction buffer supplied with the biotin tyramide kit. The reaction was performed as described for the Biotin-XX-Tyramide Superboost kit with only one modification. The reaction was incubated in an Eppendorf Thermomixer set to 25 °C with 300 rpm shaking for 10 minutes. The reaction was neutralized by adding PBS + 10mM sodium ascorbate, 10mM Trolox and 10mM sodium azide to halt HRP activity and pelleted at 200g for 5 minutes. The cells were then resuspended in PBS + 0.001% Tween-20 by flicking. A small aliquot was taken to confirm the reaction by fluorescence staining (see below) and the remainder was pelleted and stored at −80 °C or immediately used for fragmentation and Streptavidin pull down (see below).

### Fluorescence staining for the cTSA-seq reaction

Cells from the completed cTSA-seq reaction were resuspended in PBS containing 0.001% Tween-20 and Streptavidin conjugated to Alexa 488 (1:200, Biolegend, #405235) and an anti-rabbit secondary conjugated (1:1000) to an Alexa 594 fluorophore. The cells were then stained with DAPI and mounted in Prolong anti-fade gold for imaging on a Leica SP5 scanning confocal microscope using the Leica Application Suite v2.7.3.9723. We used a 40x 1.4NA objective with Leica Type F immersion oil. Diffusion of the Streptavidin signal from the lamin-B1 source signal was calculated using the iterative Levenberg-Marquardt algorithm R package minpack.lm v.1.2-1.

### Streptavidin pull down of chromatin at or near the nuclear lamina

The cells were resuspended in 300μl RIPA buffer (50 mM Tris, 150 mM NaCl, 0.1% (wt/vol) SDS, 0.5% (wt/vol) sodium deoxycholate and 1% (vol/vol) Triton X-100, pH 7.5) supplemented with phenylmethylsulfonyl fluoride (PMSF) and incubated at 4 °C with end-over-end rotation for 30 minutes. The suspension was sonicated on a Diagenode Bioruptor Pico and a 50μl aliquot of the resulting lysate was collected for the input while the remainder was used for the pull down portion of the pulldown experiment. The input was digested with 10μl of 10mg/ml Proteinase K overnight at 50 °C with 700 rpm shaking. For the Streptavidin pull down (StrePD), Streptavidin coated magnetic beads (Pierce, #88817) were washed and resuspended in RIPA buffer. We added ∼70μl suspension of Streptavidin magnetic beads per pull down. The following day, the beads were collected and washed sequentially with two times in 1ml RIPA, one time in 1ml LiCl buffer (250mM LiCl, 1% IGEPAL-CA630, 1% deoxycholic acid, 10mM Tris, pH 8.0, 1mM EDTA), one time in 1ml with high salt buffer (1M KCl, 50mM Tris-Cl pH 8.0, 5mM EDTA), one time in 1ml Urea wash buffer (2M Urea, 10mM Tris-Cl pH 8.0), and then one time in 1ml RIPA. The beads were resuspended in 50μl RIPA buffer and digested with 5μl of 10mg/ml Proteinase K overnight at 50 °C with 700 rpm shaking. The input and Streptavidin pull down (StrePD) DNA were purified using Ampure XP beads (Agencourt).

### Thiol (SH)-Linked Alkylation for the Metabolic sequencing of RNA (SLAM-seq)

To determine new transcription for FACS isolated unsynchronized G1 (from asynchronous cell culture) or synchronized early and later G1 cells, we incubated asynchronous HCT116 cultures or re-seeded mitotic shake-offs of HCT116 cells with 500μM 4-thiouridine (4sU) for 1 hour prior to harvest. Twenty minutes before harvest, we pre-stained cellular DNA by directly adding Hoechst 33342 to a final of 10μg/ml. The cells were harvested by trypsinization, pelleted at 150g for 3 minutes each and resuspended in HBSS + 2% FBS supplemented with 10μg/ml Hoechst 33342 that was pre-warmed to 37 °C. The cells were immediately filtered with a 0.45 micron filter and subject to FACS as described above with the exception that the cell sorting chamber was also set to 37 °C since the cells were alive. FACS was performed for a maximum of 10 minutes, which usually yielded anywhere between 30,000 and 100,000 cells. The collected cells were then pelleted at 500g for 3 minutes, aspirated with a P100 pipette and RNA was immediately extracted using the Qiagen RNeasy Plus kit and quantitated. To convert the incorporated 4sU, we incubated the RNA (∼0.5-1μg) in PBS pH 7.4 with 100μM iodoacetamide at 50 °C for 15 minutes and then re-purified the RNA using the Qiagen RNeasy Plus kit. As a control, we performed iodoacetamide treatment and sequencing on unlabeled (no 4sU treatment) asynchronous G1 cells to provide an estimation of sequencing errors.

### Sequencing library production

RNA library building was done using the Illumina TruSeq RNA library kit v2 with Ribo-depletion. DNA libraries were prepared using the Rubicon Genomics ThruPlex kit. Sequencing was performed on the Illumina NextSeq 500 platform.

### Data processing

Mapping of raw reads was done using the hg19 or mm9 assemblies and Bowtie 2.3.2 under the default setting. Duplicates were removed using samtools v1.6. The reads were then called into 10 or 100 kilobase (kb) genomic windows using the coverage function in bedtools v2.26.0. Next, we normalized the StrePD and input read count data to one million, calculated the log2(StrePD/input) value and transformed the data to a z-score. The publicly available LADs and lamin-B1 ChIPseq datasets examined in this study were analyzed as described above. The depmixS4 v1.4-0 R package was used to call a three-state Hidden Markov Model (HMM) to identify LADs coordinates. The three-state model parsed the cTSA-seq signal into regions that contained 1) clear NL-chromatin signal enrichment, 2) intermediate or noisy signal and 3) clear depletion. We intersected the HMM calls between experimental replicates to find LAD regions that were reproducibly identified. Quantitation of epigenetic and lamin-B1 signals was done using the bigwigAverageOverBed function from the Kentutils v3.62 package. For analyses that examined chromosomes as a percentage of their length, we first split chromosomes in 200 windows using the makewindows function in bedtools and then quantitated the lamin-B1 signal over these windows using the bigwigAverageOverBed function from Kentutils. A/B compartment calls were done using CscoreTool v1.1 (31). Analysis of sequencing depth was done with Preseq v2.0.3. Additional analysis, statistics and graphical plotting were done either in R Studio v0.98.95.3 using the pHeatmap v1.0.12, corrplot v0.84 and Hmisc v4.2-0 packages or Microsoft Excel 2016. Browser tracks were displayed using the UCSC Genome Browser with a smoothing window of 2.

For SLAM-seq experiments reads were initially trimmed by 10 nucleotides from both the 5’ and 3’ ends using cutadapt v1.6. The trimmed RNA-seq data was aligned with Bowtie 2.3.2 and Tophat2 under default settings using the Ensembl hs75 assembly. Polymorphisms corresponding to the expected U- to C-conversions were quantitated using the GrandSLAM v 2.0.5c package (32). Data was filtered for a minimum of 10 transcripts per million (TPM) and the new to total RNA (NTR, “MAP”) value was used to represent the gene for subsequent analysis. Statistics and graphical plotting were done either in R Studio v0.98.95.3 using base functions, pHeatmap v1.0.12, corrplot v0.84, car v3.0-5 and Hmisc v4.2-0 packages, or Microsoft Excel 2016.

### Statistical analyses

Pearson correlations seen in Figures 1D, 1F, 3B and S2D were calculated using the rcorr(x, method=”pearson”) function from the Hmisc v4.2-0 package or the cor.test function in base R. Two-sided t-tests calculated for Figures 4D and S4E were calculated using the base R function t.test().

**Figure 1:**
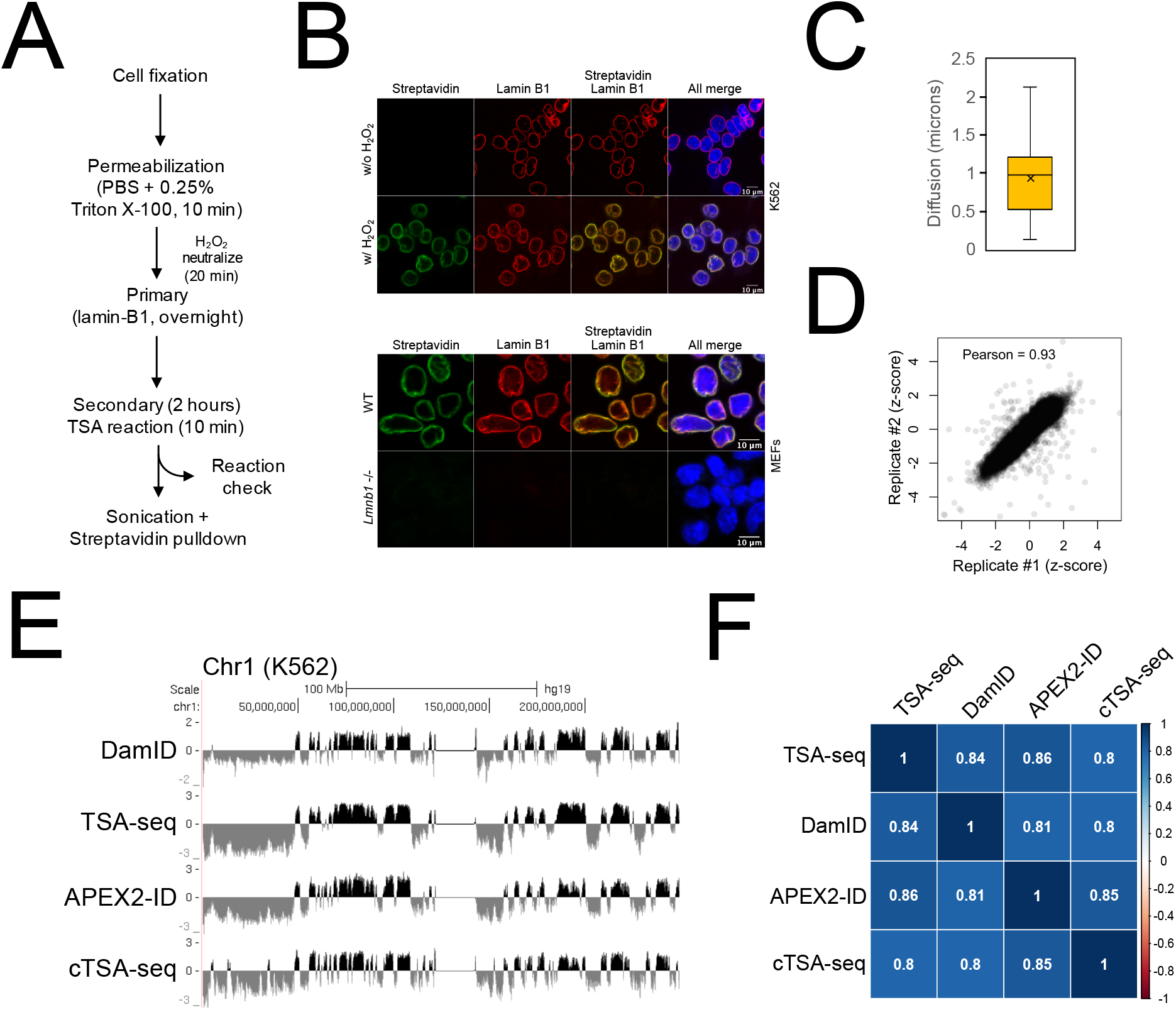
The cTSA-seq method maps chromatin at and near the NL. **(A)** The cTSA-seq experimental pipeline used in this study. **(B)** The lamin-B1 cTSA-seq reaction done with or without hydrogen peroxide using K562 cells (top panel) and with hydrogen peroxide in control (“WT”) and *Lmnb1* -/- MEFs (bottom panel). Staining was done with Streptavidin (green) to highlight biotinylated material and a secondary antibody toward the lamin-B1 antibody (red). Scale bar equals 10 microns. **(C)** Boxplot showing the observed distance of diffusion (in microns) of the cTSA-seq labeling reaction from the lamin-B1 source. **(D)** Scatterplot showing the relationship between the cTSA-seq replicates. The Pearson coefficient is 0.93. **(E)** UCSC Genome browser view of lamin-B1 cTSA-seq mapping profile in the K562 cells (Chr1, hg19). Previously published APEX2-ID, DamID and TSA-seq mapping profiles from K562 cells are shown for comparison. The y-axis represents an averaged z-score for each dataset. (**F**) Pearson correlation matrix for DamID, TSA-seq, APEX2-ID and cTSA-seq.

### Data Availability/Sequence Data Resources

Sequencing data generated for this study is deposited at NCBI GEO Accession number GSE186503. K562 cell DamID and TSA-seq LADs data (21) was obtained from Gene Expression Omnibus (GEO) GSE66019. The K562 cell APEX2-ID data (8) was from GEO GSE159482. K562 ENCODE RNA-seq data was obtained from GEO GSM958731. The K562 ENCODE ChIPseq data was obtained from GEO GSM733776. The Tig3 OIS data (29) was obtained from GEO SE76605 and the IMR90 OIS data (30) was obtained from GEO GSE49341. The epigenome data for HCT116 cells was obtained from the ENCODE project (HCT116 reference epigenome series ENCSR361KMF). The HCT116 ENCODE RNAseq data set was obtained from GEO GSM958749. The K562 Hi-C dataset (4) was obtained from GEO GSE63525, and the SNIPER K562 subcompartment track was obtained from https://cmu.app.box.com/s/n4jh3utmitzl88264s8bzsfcjhqnhaa0/folder/86847304302 (33). UCSC Genome browser tracks are available at the following URLs:

Figures 1E, 2A and S1D:

**Figure 2:**
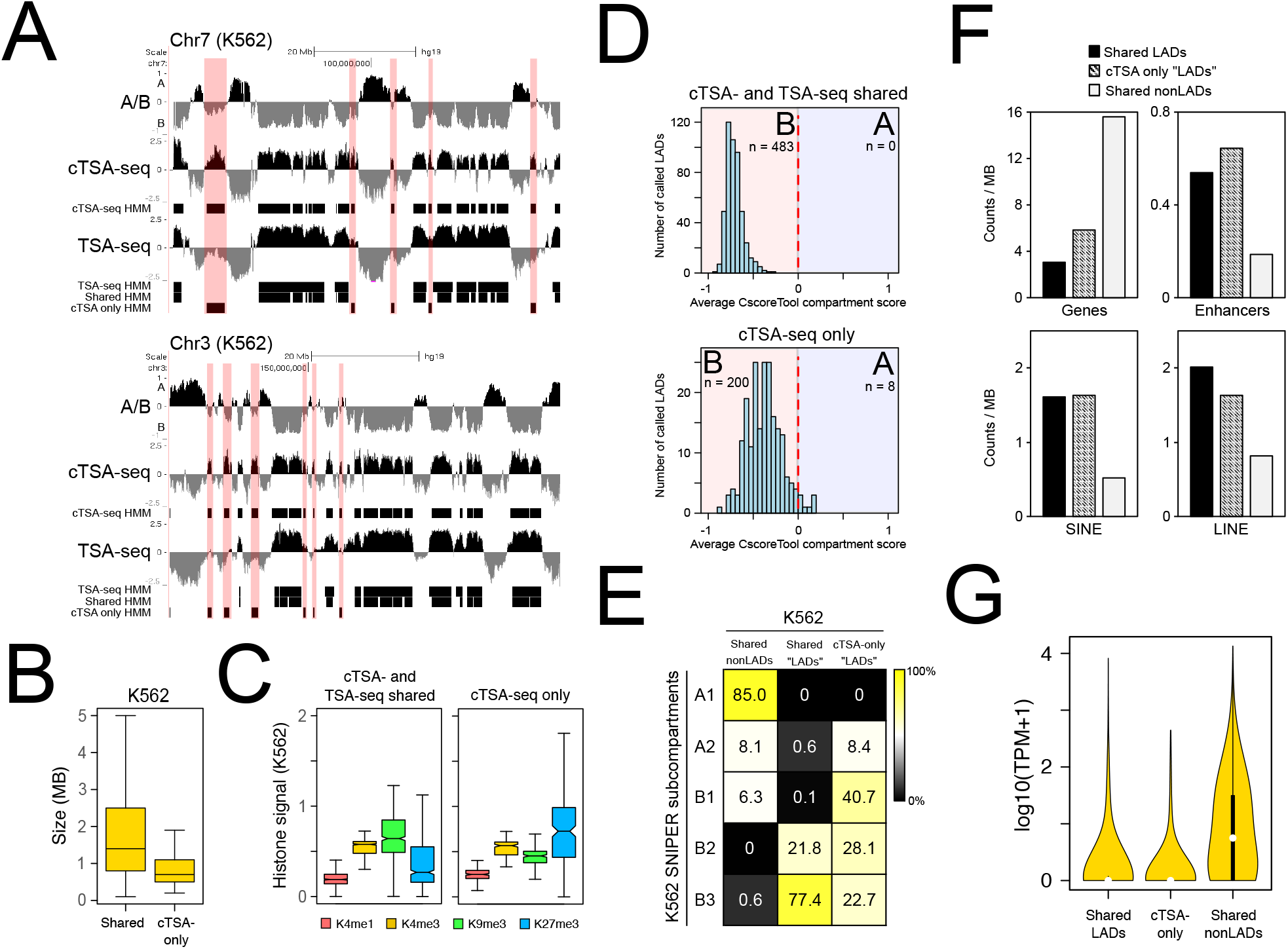
The cTSA-seq identified chromatin regions are present in the B compartment. **(A)** Example UCSC Genome Browser tracks for select regions of chromosome 7 and 3 (hg19). The K562 Hi-C dataset is presented as an A/B compartment track (“A/B”) where the A compartment is represented by positive values and the B compartment by negative values. cTSA-seq and TSA-seq data is presented as an averaged z-score. The three-state HMM tracks are shown below each track. The “Shared HMM” track represents an intersection between cTSA-seq and TSA-seq HMM calls. The light red bars highlight regions that were called by the HMM as strong signal for cTSA-seq only and coincides with the “cTSA-seq only HMM” tracks. **(B)** Boxplot showing the size of cTSA-seq and TSA-seq (“Shared”) NL-chromatin regions and cTSA-seq only (“cTSA-only”) regions. **(C)** Boxplots showing the histone modification signals for “cTSA- and TSA-seq shared” NL-chromatin regions (left) and “cTSA-seq only” regions (right). H3K4me1 is shown in red, H3K4me3 in yellow, H3K9me3 in green and H3K27me3 in blue. **(D)** Frequency histograms showing the number of called chromatin regions at or near NL, referred to as “LADs”, and their average compartment score. The top panel pertains to the “cTSA- and TSA-seq shared” regions while the bottom is related to “cTSA-seq only” regions. The B-compartment is shaded light red and the A compartment is shaded blue. The number of called “LADs” falling into each compartment is noted beneath the “A” or “B” label for each compartment. **(E)** Heatmap showing the percentage of SNIPER-defined K562 Hi-C subcompartments in K562 shared cTSA- and TSA-seq (“Shared LADs”) or cTSA-seq only chromatin regions (“CTSA-only LADs”). Shared nonLADs refers to euchromatic regions identified by both methods. **(F)** Bar charts showing the number of genes (upper left), enhancers (upper right), SINE elements (lower left) and LINE elements (lower right) in LADs shared between cTSA- and TSA-seq (“Shared LADs), cTSA-seq only LADs (“cTSA-only”) and in “Shared nonLADs” regions. **(G)** Violin plot showing the expression level of genes found in “Shared LADs”, cTSA-seq only “LADs” (“cTSA-only”) and in “Shared nonLADs” regions.

https://genome.ucsc.edu/s/tranjoseph/hg19_K562_Figures_1%262

Figure 3A:

**Figure 3:**
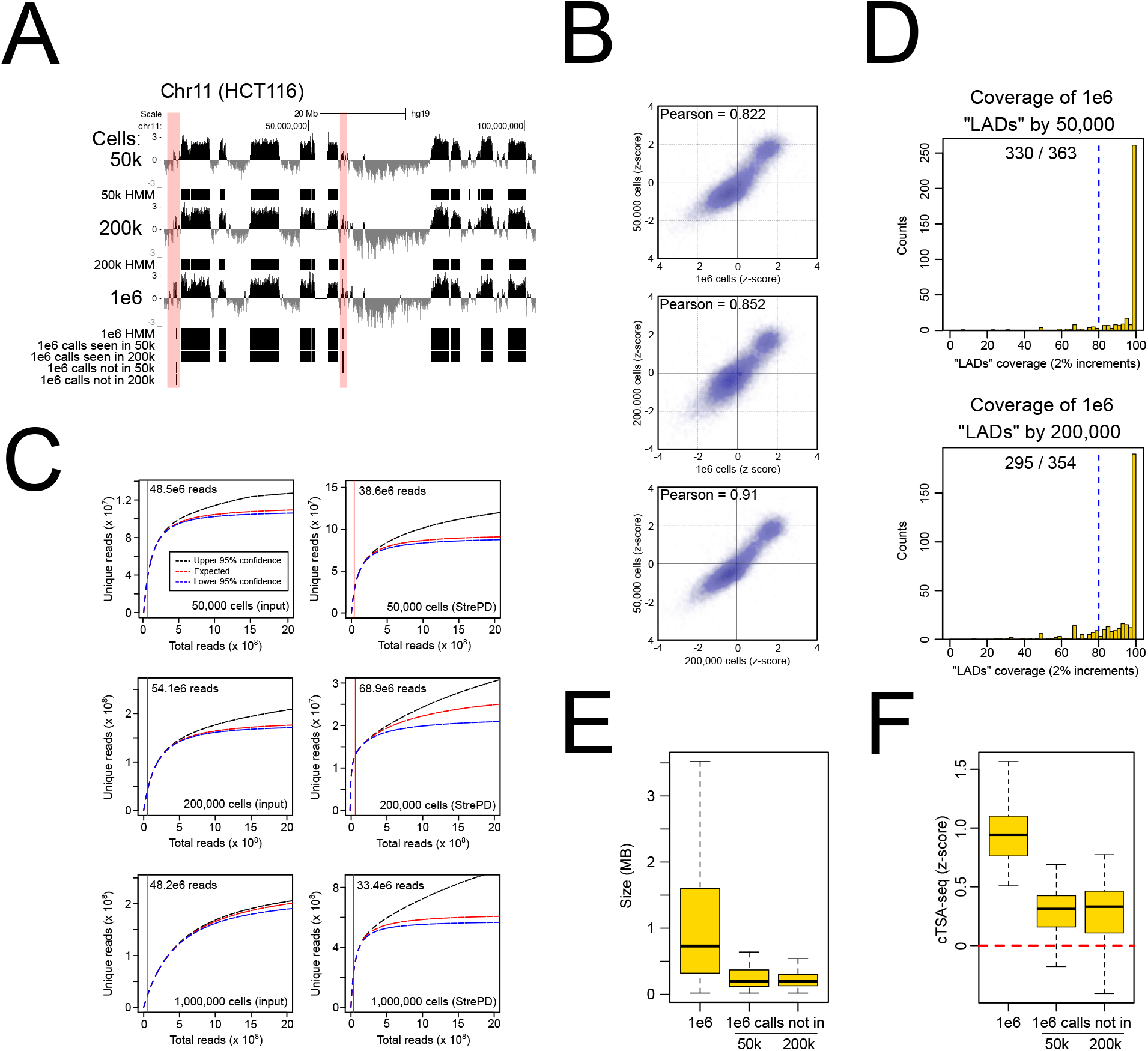
cTSA-seq can map chromatin at or near the NL from as little as 50,000 cells. **(A)** UCSC Genome Browser tracks for low cell cTSA-seq (Chr11, hg19). “50k”, “200k” and “1e6” represent the 50,000, 200,000 and 1,000,000 HCT116 cells, respectively. Three-state HMM tracks, corresponding to the strongest signals, are shown beneath each signal track. 1e6 HMM calls identified (“1e6 calls seen in 50k” and “1e6 calls seen in 200k”) and not identified (“1e6 calls not in 50k” and “1e6 calls not in 200k”) by low cell experiments are shown at the bottom. Light red bars highlight regions from the one million cell sample not called or called as weak signal by the HMM using data from low cell experiments. **(B)** Scatterplots comparing data (z-score) from 50,000, 200,000 and 1,000,000 cell cTSA-seq experiments. Pearson correlation coefficient is shown on the upper left of each plot. **(C)** Representative analysis of sequencing depth and the predicted number of unique reads for each sample from the 50,000, 200,000 and 1,000,000 cell cTSA-seq. The red vertical line indicates the actual sequencing depth for each sample. The read numbers are shown labeled on the upper left. Input samples are the left three plots while Streptavidin pull down (“StrePD”) samples are on the right and these are denoted on the lower right. **(D)** The coverage of HMM-defined “LADs” regions from the one million cell dataset by those obtained from the low-cell experiments. The coverage of the one million cell by the 50,000 cell data is the top graph while the coverage by 200,000 cells is the bottom graph. The data is presented as a histogram where the y-axis is the number of “LADs” presented as “counts” and the x-axis is the percentage of the “LAD” covered. The coverage bins are in 2% increments. The blue dashed line is the 80% coverage position. The actual number of “LADs” with 80% or greater coverage over the total number of LADs is shown in the middle top part of each graph. **(E)** Boxplot showing the size (in megabases) of the mapped chromatin regions from 1,000,000 cells, and those from the 1,000,000 cell experiment not called by the lower cell number experiments. **(F)** Boxplot showing the cTSA-seq (z-score) signal of HMM-defined chromatin regions from 1,000,000 cells, and those from the 1,000,000 cell experiment that were not called in the lower cell number experiments.

https://genome.ucsc.edu/s/tranjoseph/hg19_low_cell_HCT116_cTSAseq_Figure_3

Figure 5A, 5C, 5F, S4A, S4C:

https://genome.ucsc.edu/s/tranjoseph/hg19_R90_R180_G1SG2_Lenain_Sadaie_RPE_FIgure_5%2 6S4

Figure S3B:

https://genome.ucsc.edu/s/tranjoseph/hg19_R90_R180_Async_G1_rank_check_Figure_S3

## Results and Discussion

### Development of chromatin pull down-based Tyramide Signal Amplification-sequencing (cTSA-seq) to reliably map chromatin regions at or near the NL

The Tyramide Signal Amplification (TSA)-seq method utilizes the small fraction of DNA that is labeled by the TSA reaction to map LADs (21). However, the TSA labeling occurs on both DNA and proteins (21,25) that are at or near the nuclear lamina. We first examined the Tyramide Signal Amplification (TSA) reaction with a commercial kit (see Methods) using K562, HCT116, and mouse embryonic fibroblast (MEF) cell lines and a commercially available rabbit lamin-B1 antibody (Abcam, #ab16048) to target the proximity labeling of proteins at or near the NL with biotin (Figure 1A). The post-reaction staining with an Alexa-conjugated Streptavidin coincided largely with lamin-B1 while Streptavidin staining was not evident in control reactions that omitted the hydrogen peroxide (Figure 1B, top panel), used an anti-mouse HRP secondary antibody (Figure S1A) or used lamin-B1 null MEFs (Figure 1B, bottom panel). Similar to previous work (21), the diffusion of Streptavidin signal for our unmodified TSA reaction was ∼1 μm from the primary lamin-B1 antibody label (Figure 1C).

Next, we sonicated and pulled down the fragmented biotinylated chromatin (Figure S1B) with Streptavidin beads for sequencing (Figure 1A). We used one million K562 cells, which is at least 10 times less than that required for TSA-seq. The data obtained from cTSA-seq biological replicates were highly correlated (Figure 1D, Pearson = 0.93) and the average cTSA-seq signal was similar to those obtained by DamID, the original TSA-seq method and our recently adapted APEX2-ID method (Figure 1E, F). To further assess the cTSA-seq data, we performed decile analysis as done for SON TSA-seq (21). The published decile analysis for SON TSA-seq data revealed a gradient of epigenetic signal and transcription activity across the nuclear speckle-lamin TSA-seq axis (21), with elevated levels of transcriptional activity and euchromatic marks closest to the nuclear speckle. Conversely, these signatures were weakest at the NL. We stratified the lamin-B1 cTSA-seq signal into decile ranks where the 1st and 10th ranks corresponded to the lowest and highest cTSA-seq log2 values, respectively (Figure S1C, S1D); and then measured the TSA-seq, epigenetic and transcriptional signal within genomic regions defined by each decile. Similar to what was observed for TSA-seq, cTSA-seq deciles showed an increasing gradient of transcriptional activity and euchromatic epigenetic signal away from the NL, which is represented by the lower deciles (Figure S1E). As anticipated, elevated heterochromatic signatures were closest to the NL (Figure S1E).

We next used a Hidden Markov Model (HMM) to identify the NL-chromatin regions and then intersected the calls from each replicate to identify the strongest signals that were also reproducible. This approach has been previously used to identify NL-associated chromatin with the APEX2-ID (8) and pA-DamID genome mapping methods (7). The three-state HMM model used here, however, separates the cTSA-seq data into 1) strong signal representing clear enrichment, 2) intermediate or noisy signal and 3) signal that is not enriched by the procedure. The HMM calls contain the highest decile ranking signal (Figure S1D, compare “cTSA-seq HMM” and ranks 7-10 tracks). Based on this, we found that the cTSA-seq approach identified 97.3% of the LADs identified by the same three-state HMM model from data generated by the related TSA-seq method (21).

Interestingly, the HMM identified cTSA-seq-specific enrichments that were not called from the TSA-seq data, but visually most appeared to coincide with regions that showed weak enrichment or depletion of TSA-seq signal (Figure 2A, light red bars). These lamin-B1 cTSA-seq-specific chromatin regions were often centered around a log2 of 0 in the TSA-seq data (Figure S1F) and indicated that these regions were frequently neither enriched nor depleted by the TSA-seq method. These cTSA-seq specific regions were generally smaller in size (Figure 2B, <1MB) but were enriched for H3K27me3 (Figure 2C), which suggested that these were heterochromatic regions.

To gain further understanding of the cTSA-seq specific regions, we examined the Hi-C compartments associated with these chromatin regions. Previous analyses of Hi-C data have broadly identified the A and B compartments, which correspond to open euchromatin and closed heterochromatin, respectively (3,4). As anticipated, all chromatin regions identified by both lamin-B TSA- and lamin-B1 cTSA-seq methods (“shared” regions) were enriched for B-compartment scores calculated from the K562 Hi-C dataset by the CscoreTool program developed in our lab (Figure 2D, top panel) (4,31). Most (96.2%, 200 out of 208) of the cTSA-seq-specific chromatin regions were also classified as the B-compartment (Figure 2D, bottom panel). To further explore this observation, we compared our cTSA-seq data with the published SNIPER K562 subcompartments track (4,33). The SNIPER algorithm reliably subdivides the A and B compartments in the A1-2 and B1-B3 subcompartments, respectively, but excludes the B4 subcompartment since this subcompartment was only found on chromosome 19 (33). The TSA- and cTSA-seq shared chromatin regions, which we referred to as “LADs”, are enriched for B3 and B2 compartments (99.2%), while cTSA-seq specific regions contain signatures for B1, B2 and B3 compartments (91.5%), with a notable bias toward the B1 compartment (Figure 2E). We further found that cTSA-seq specific regions are more enriched for genes, enhancers, SINE elements and contained lower LINE element content than TSA- and cTSA-seq shared “LADs” regions (Figure 2F). Both the shared and cTSA-seq specific regions have low gene expression (Figure 2G). These analyses show the cTSA-seq method can be used to map chromatin in the B-compartments that includes previously defined LADs and heterochromatin regions that are near the NL.

The identification of chromatin regions found in the B-compartment but not in the previously defined LADs is interesting as it suggests that the B-compartment defined by Hi-C contains both LADs and chromatin regions near LADs. The ease of biotinylation of proteins near the NL by diffusion may lead to the pull down of chromatin proximal to the NL that is in a closed chromatin state. Additionally, the chromatin based pull down by the cTSA-seq method may capture both labeled naked DNA and chromatinized DNA thereby increasing mapping sensitivity.

### cTSA-seq mapping of chromatin at or near the NL in as few as 50,000 cells without adding carriers

The efficient mapping of chromatin regions in LADs or near the NL from low numbers of cells would open up the possibility of characterizing small number of pure cells directly isolated from *in vivo* sources without further culturing or from subpopulations of cells sorted from *in vitro* culture. Thus far, no LADs mapping has been done for low-cell populations, except for one report of single cell LADs mapping by DamID, which suffers from data sparsity (19).

We examined the possible use of cTSA-seq to map chromatin regions at or near the NL in a lower number of cells. We titrated HCT116 cells down to 50,000 and 200,000 cells prior to performing immunostaining with Lamin-B1 and cTSA-seq. A one-tenth input was collected from the sonicated lysate prior to Streptavidin pull down. The lamin-B1 cTSA-seq profiles from the 50,000 and 200,000 cell samples visually appeared similar to that mapped from one million cells, indicating that cTSA-seq method enables high quality mapping of chromatin at or near NL in the B-compartment using as few as 50,000 cells (Figure 3A). The 50,000-cell dataset was well correlated with both the 200,000 and 1,000,000 cell datasets (Pearson >0.82, Figure 3B). Based on the unique and total reads, our sequencing depth did not reach saturation (Figure 3C) and we were able to achieve good mapping from as few as 33.4 million reads.

We next applied the same three-state HMM model used above to our low-cell data to call the strongest reproducible chromatin regions at or near the NL (Figure 3A, “HMM” tracks). Intersecting these HMM calls revealed that the low-cell cTSA-seq can identify the majority of NL-chromatin regions (>82%) found in the 1 million cell reference sample. Most of these 1 million cells reference “LADs” also identified by the low cell data (84-91%) showed 80% or greater coverage by the low cell mapping (Figure 3D), and indicates that we can achieve reasonably complete mapping from 50,000 cells. We did notice that smaller regions with an average size of ∼250kb were less efficiently identified by the HMM model when using data from low-cell cTSA-seq experiments (Figure 3A, light red boxes, Figure 3E). However, these regions missed by the HMM calls had relatively weak signals in all samples (Figure 3F) suggesting that these missed calls were filtered out by the three-state HMM model.

We show that cTSA-seq can offer high quality mapping of chromatin at or near the NL in as few as 50,000 cells without adding any carriers. We have previously developed a carrier approach to reduce sample loss in low-cell ChIP-seq, and have successfully used this approach to map epigenetic marks and transcription factor binding sites from as few as 100 cells (34,35). Applying this carrier approach to cTSA-seq should enable mapping of chromatin regions at or near the NL in far fewer cells than 50,000, especially considering that these chromatin regions represents 30-50% of the genome. Overall, the data presented here shows that cTSA-seq is a complementary tool that can efficiently identify chromatin regions consisting of dominantly LADs and NL-proximal chromatin found in the B compartment from as few as 50,000 cells.

### cTSA-seq enables the use of sorted cells to more accurately identify chromatin regions at or near the assembling NL as cells progress through G1 phase

A recent study on the dynamics of LADs during the cell cycle revealed an increased interaction between the distal ends of chromosomes and the reassembling NL during early G1 in synchronized cell populations (7). This study collected all cells at a fixed time point after mitotic block and release for mapping. However, mitotic block does not arrest all cells in mitosis and the arrested cells do not enter interphase with perfect synchrony after release (Figure S2A). Therefore, mapping using all released cells would affect the results. Since cTSA-seq offers high quality mapping of only 50,000 cells, we reasoned that we should be able to improve the mapping by adding a sorting step for G1 DNA content to exclude contaminating un-arrested cells or cells that have not entered into G1.

We performed nocodazole arrest of HCT116 cells followed by a mitotic shake-off and release into fresh media (Figure 4A). Fluorescence Activated Cell Sorting (FACS) analyses showed that some mitotic HCT116 cells released from nocodazole progressed into G1 of the cell cycle (Figure S2A, left panel). A clear but minor G1 cell peak could be seen between 60-90 minutes after release, and by 180-240 minutes the profile was similar to that seen for asynchronous cultures suggesting that much of the cell population had finished mitosis (Figure S2A, left panel). We next used this approach to FACS isolate the G1 cell population at 90 or 180 minutes after mitotic release. These cell populations will be referred to as “90m G1” or “180m G1” cells, respectively, to differentiate them from the G1 population isolated from asynchronous cell populations (“Async G1”). We found that both 90m and 180m G1 cells have entered G1 as judged by DAPI and lamin-B1 staining (Figure 4B). Importantly, lamin-B1 staining appeared strongly at the nuclear periphery thus allowing the use of cTSA-seq. Chromatin in many 90m G1 nuclei appeared noticeably dissimilar to 180m G1 and asynchronously growing G1 HCT116 cells (Figure S2B) as has been previously observed for synchronized cells released from mitotic block (36). Using the same nocodazole block and release protocol, we similarly isolated early (90m) or later (180m) G1 hTERT-RPE1 cells. Based on the FACS profile, hTERT-RPE1 cells progressed into G1 slower (7) than that of HCT116 cells (Figure S2A, right panels), but similar to HCT116 cells, staining revealed that cells have entered G1 at both time points (Figure S2C).

**Figure 4:**
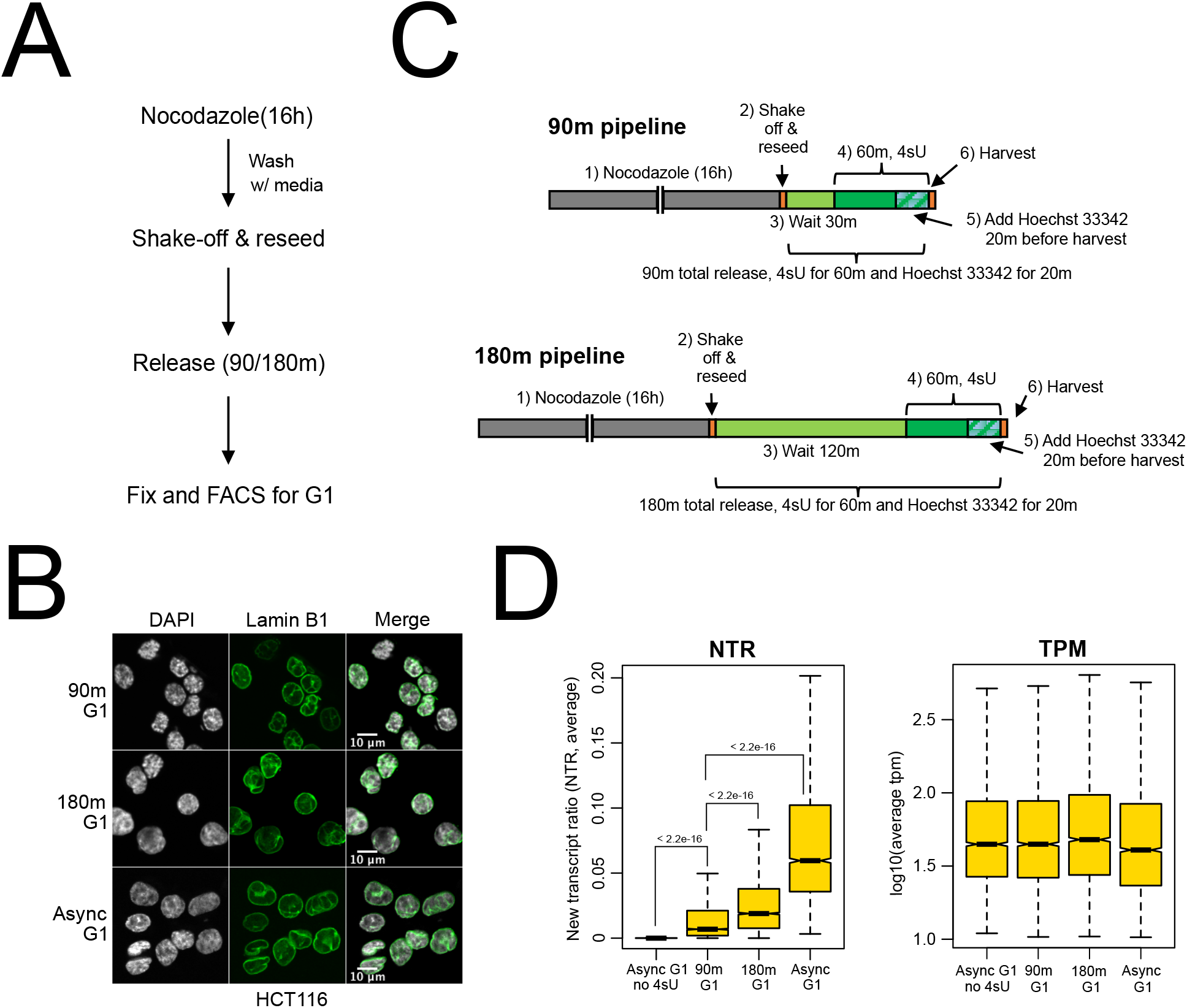
Characterization of early and later G1 cells. **(A)** Illustration of experimental workflow for nocodazole block and release protocol. **(B)** Lamin-B1 immunostaining (green) for HCT116 cells in early G1 (“90m G1”), later G1 (“180m G1”) and asynchronous G1 (“Async G1”) isolated from asynchronous cell cultures. DAPI is presented in gray and lamin-B1 is presented in green. The scale bar equals 10 microns. **(C)** Experimental pipeline for performing SLAM-seq experiments on early (“90m pipeline”) and later (“180m pipeline”) G1 HCT116 cells. The 4sU labeling time was kept constant at 60 minutes for each sample. **(D)** Boxplots showing the average new transcripts to all transcripts ratio (NTR, left) and a log10 transformation of the transcripts per million (TPM, right) for indicated samples. The p-values shown in the boxplot were calculated from a two-tailed t-test.

Next, we further confirmed that the 90m and 180m G1 cells described above represented the early and later stages of G1, respectively. Previous reports indicate that transcription progressively increases in post-mitotic G1 (37,38). To accomplish this, we used the Thiol (SH)-Linked Alkylation for the Metabolic sequencing of RNA (SLAM-seq) labeling strategy (Figure 4C) to measure new transcripts (39). Cells treated with 4-thiouridine (4sU) incorporate the analog into newly transcribed RNA (40), and treatment of RNA with iodoacetamide results in a U- to C-conversion of the 4sU analog during reverse transcription that can be quantitated and expressed as a ratio of new RNA transcripts to total RNA (NTR) (39,41). We performed SLAM-seq on 90m G1, 180m G1, and asynchronous G1 samples by labeling with 4sU for 60 minutes prior to the harvesting time point (Figure 4C). The replicate experiments were concordant (Figure S2D), and a low level of new transcription was observed in 90m G1 cells that further increased in the 180m G1 cells (Figure 4D, left panel, p-value < 2.2e^-16^) even though global transcript levels remained similar (Figure 4D, right panel). The expression of select cell cycle related (e.g., *CDT1, CDK4, CDK6*, and *PCNA*) and DNA replication related genes (e.g., *ORC1, ORC6, MCM3,7,8*) were not prominent in 90m G1 samples, but was more apparent in 180m G1 and asynchronous G1 samples (Figure S2E). These results are consistent with previous observations and show that the 90m G1 fraction contained cells that has entered early G1 phase characterized by low levels of transcription, while the 180m G1 cells represent a later stage of G1 with an increased number of genes being expressed.

We next used cTSA-seq to identify chromatin regions at or near the reassembling NL in early and later G1 phase HCT116 cells. We also performed cTSA-seq of asynchronous G1, S, and G2 cells isolated by FACS, which were previously shown to be largely similar with minor variations (7,8). The G2/M population was referred to as “G2” since the nuclear envelope is disassembled during mitosis and does not contribute much to the signal (19,42). Lamin-B1 cTSA-seq mapping revealed a graded enrichment of signals toward chromosome ends in 90m G1 HCT116 cells, suggesting that the reassembling NL is closest to these chromosomal regions (Figure 5A, 5B grey bars). cTSA-seq mapping of the later 180m G1 population revealed a signal reduction of chromosome ends and the appearance of chromatin regions that were mapped in the asynchronous G1, S and G2 populations (Figure 5A, 5B, Pearson > 0.7). We also performed the same experiment with hTERT-RPE1 and found a similar enrichment of signal toward the distal ends of chromosomes at the 90m early (Figure 5C, 5D). Our lamin-B1 cTSA-seq mapping pattern of the early 90m G1 cells is similar to the early NL-chromatin interactions previously observed using an antibody-targeted Dam-ID approach (7). Thus, the cTSA-seq method readily identifies chromatin at or near the reassembling NL in early G1 from FACS sorted cell populations.

**Figure 5:**
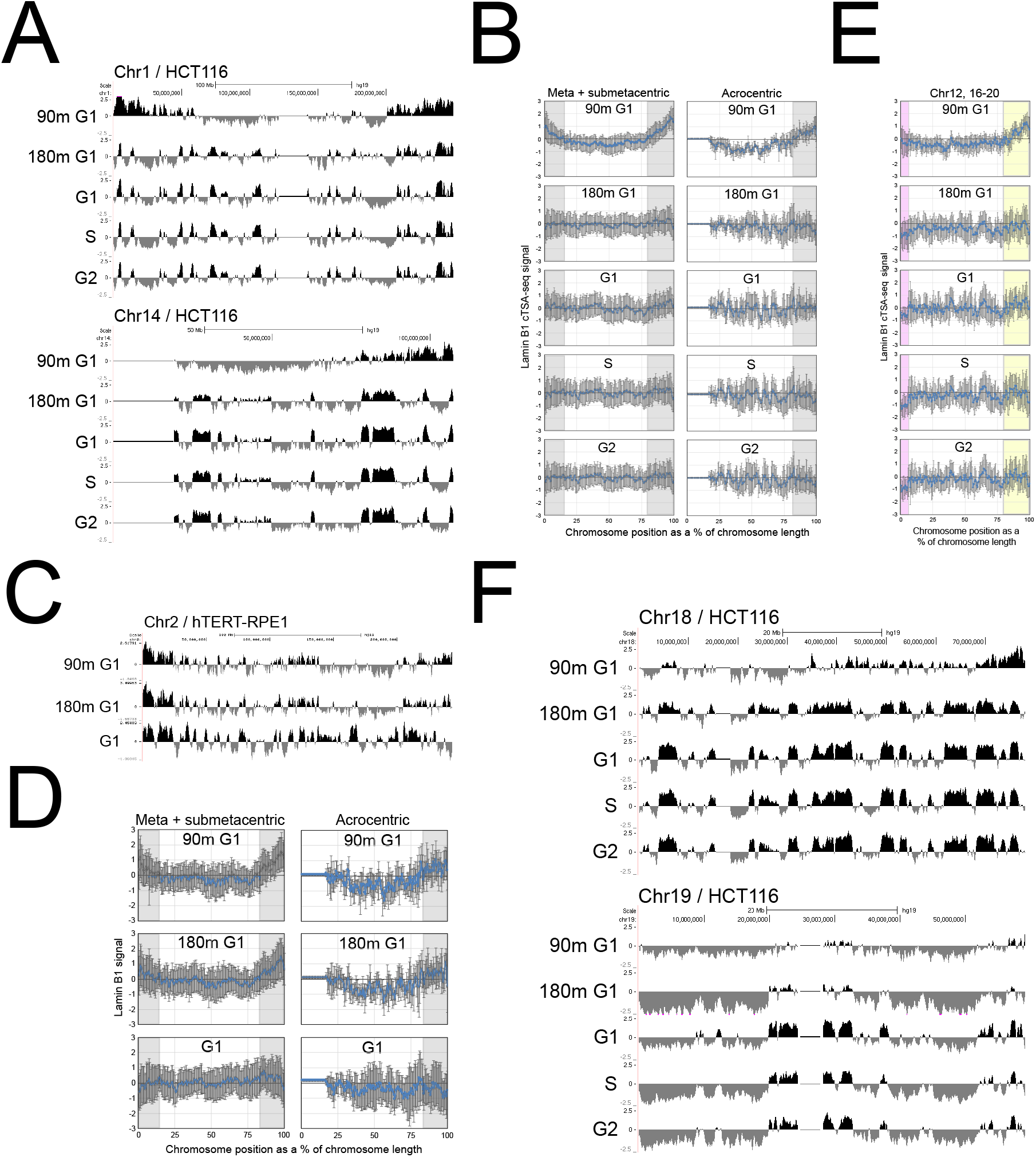
The cTSA-seq identifies early G1 chromatin interacting with or near the reassembling NL. **(A)** UCSC Genome browser tracks (hg19) showing the signal obtained by lamin-B1 cTSA-seq from early G1 (“90m G1”), later G1 (“180m G1”), asynchronous G1 (“G1”), S and G2 sorted HCT116 cells. The metacentric chromosome 1 (top) and acrocentric chromosome 14 (bottom) are displayed. The data presented are average z-scores of both replicate experiments. **(B)** Lineplots showing the average lamin-B1 cTSA-seq signal from meta- and submetacentric (left) and acrocentric (right) chromosomes in the indicated cell cycle stages. Each chromosome was segmented as a percentage of their length and the lamin-B1 signal was quantified over these windows. Error bars represent standard deviation of signal. Grey bars highlight distal regions of the chromosome that show increased cTSA-seq signal in early G1. **(C)** UCSC Genome Browser tracks (Chr2 2, hg19) mapped by lamin-B1 cTSA-seq from “90m G1”, “180m G1” and asynchronous G1 (“G1”) populations of hTERT-RPE1 cells. **(D)** Average lamin-B1 cTSA-seq signal from meta- and submetacentric (left) and acrocentric (right) hTERT-RPE1 chromosomes. Each chromosome was evenly segmented as a percentage of their length and the cTSA-seq signal was quantified over these windows. The error bars represent standard deviation. Grey bars highlight regions where cTSA-seq signal is enriched during early G1. **(E)** The average lamin-B1 cTSA-seq signal for smaller (chromosomes 12, 16-20) meta- and submetacentric chromosomes. Purple bars highlight the low cTSA-seq signal seen on the short arm of these chromosomes that further decreases in the other samples. The yellow bars highlight the stronger distal cTSA-seq signal seen in early G1. **(F)** UCSC genome browser tracks of lamin-B1 cTSA-seq signal for chromosomes 18 (top) and 19 (bottom) across different stages of the cell cycle. The data presented are average z-scores.

In general, the distal enrichment of lamin-B1 signal revealed by cTSA-seq in early G1 cells occurred across metacentric, sub-metacentric and acrocentric chromosomes (Figure 5B, grey bars). It is notable that smaller, non-acrocentric chromosomes, such as chromosomes 12,16-20, had early G1 cTSA-seq signal that was mainly found only on the long (q) arm of the chromosome (Figure 5E, yellow bars) while the short (p) arm of these chromosomes, which showed little or no initial lamin-B1 chromatin signal, underwent a further reduction of the signal in later G1 and asynchronous G1 cells (Figure 5E, purple bars). The well-studied human chromosomes 18 and 19 are known to preferentially localize to the nuclear periphery and nucleoplasm, respectively, and their differential localization was reported to occur in early G1 (16). We found that in early G1, chromosome 18 shows an accumulation of lamin-B1 cTSA-seq signal at one end of the chromosome while chromosome 19 shows little to no distal signal (Figure 5F).

We examined for the possibility of any transcriptional effect due to these NL-chromatin dynamics. To accomplish this, we first defined NL-chromatin regions by the three-state HMM model, as done for the K562 cTSA-seq data described above. The HMM model identified regions that were based largely on signal strength and it identified similar regions as those defined by decile ranking of cTSA-seq signal (Figure S3A, S3B). The number of genes (∼5000) in chromatin regions that are preferentially near or interacting with lamin-B1 based on cTSA-seq in the 90m early G1 cells was over twice the number of those (∼2000) seen in later 180m G1, and the asynchronous G1, S and G2 cells (Figure S3C). However, we did not observe the transcription of genes, as assessed from SLAM-seq, that were transiently seen in non-LAD regions in early G1 before returning to LADs-like regions in later and asynchronous G1. This suggests that transient NL-chromatin interactions have little to no effect on gene expression in early G1.

As previously described (7,8), the vast majority of LADs mapped by cTSA-seq were stable across the G1, S and G2 stages of the cell cycle. However, minor variations in LADs across different cell cycle stages were apparent in our cTSA-seq data, as previously described in studies of cell cycle LADs using the APEX2-ID (8) and pA-DamID methods (7). These minor cell cycle-associated variable LADs (“variable”) were H3K27me3 enriched and low in lamin-B1 signal while “shared” LADs which were stable across the cell cycle were high in both H3K9me3 and lamin-B1 signal (Figure S3D). Thus, cTSA-seq confirms the observation of enriched distal chromosome interaction at or near the NL in early G1, and extends this to the HCT116 cells.

### The similarity of NL-chromatin relationships between early G1 cells and Oncogene-induced Senescence cells

The strong enrichment of lamin-B1 cTSA-seq signals toward chromosome ends in two different cell types in early G1 phase promoted us to look for similar patterns in all reported maps of NL-chromatin interactions. We noticed that LADs mapped by DamID in Oncogene-Induced Senescence (OIS) Tig3 human fibroblasts (29) showed a bias toward chromosome ends similar to what we observed in our lamin-B1 cTSA-seq in 90m early G1 cells (Figure S4A, S4B). Similarly, a second study examining RAS^V12^-induced OIS in IMR90 cells also noted a reduction of lamin-B1 chromatin immunoprecipitation (ChIP)-seq signal in the central region of chromosomes (Figure S4C, S4D) (30). The signal at the distal ends of chromosomes was significantly different in early G1 (90m G1) data when compared to both later G1 (180m G1) and asynchronous G1 samples, and between the control and OIS samples from the senescence studies (Figure S4E). It is tempting to speculate that these OIS cells were arrested at a relatively early G1 pattern of NL-chromatin relationship.

Here, we adapted a protocol based on TSA-seq, which we call cTSA-seq and demonstrate the ability of this method to map chromatin regions found in the B compartment of the genome that are at or near the NL in different tissue culture cells. We found that the cTSA-seq method maps most LADs (>97%) identified by TSA-seq but offers increased sensitivity by also detecting nuclear periphery heterochromatic regions that fall within the B-compartment defined by Hi-C. Importantly, the cTSA-seq method is capable of mapping these chromatin regions from as few as 50,000 cells, and with further technical adaptation such as the use of carriers, should be useful for much lower cell numbers such as those found *in vivo*. As a proof of principle, we applied cTSA-seq to map the previously observed dynamic proximal NL-chromatin interactions that appear during the transient early G1 stage of the cell cycle (7). The cTSA-seq method confirms the early G1 enrichment of distal ends of chromosome regions interacting with or proximal to the NL in hTERT-RPE1 cells and expands the observation to the HCT116 cell line, suggesting that this is a common feature of early G1. We note that this enrichment of proximal NL-chromatin signal at the distal ends of chromosomes in early G1 is reminiscent of those seen in OIS. The cTSA-seq method presented here is a convenient tool that can enable mapping of nuclear peripheral chromatin from low numbers of cells.

## Data Availability

All sequencing data is available at GEO GSE186503. UCSC Genome Browser tracks are available at the following links:

Figures 1E and 2A:

https://genome.ucsc.edu/s/tranjoseph/hg19_K562_Figures_1%262

Figure 3A:

https://genome.ucsc.edu/s/tranjoseph/hg19_low_cell_HCT116_cTSAseq_Figure_3

Figure 5A, 5C, 5F, S4A, S4C:

https://genome.ucsc.edu/s/tranjoseph/hg19_R90_R180_G1SG2_Lenain_Sadaie_RPE_FIgure_5%2 6S4

Figure S3B:

https://genome.ucsc.edu/s/tranjoseph/hg19_R90_R180_Async_G1_rank_check_Figure_S3

## Acknowledgements

We thank members of the Zheng and Goldman labs for advice and discussions. We also thank Allison Pinder and Frederick Tan for their help with sequencing and data analysis.

## Author contributions

J.R. Tran designed, performed and interpreted experiments. S.A. Adam, R.D. Goldman and Y. Zheng participated in the design of the study. R.D. Goldman and Y. Zheng were responsible for project funding. J.R. Tran and Y. Zheng co-wrote the manuscript. All authors participated in revisions of the manuscript.

## Figure captions

**Figure S1:**
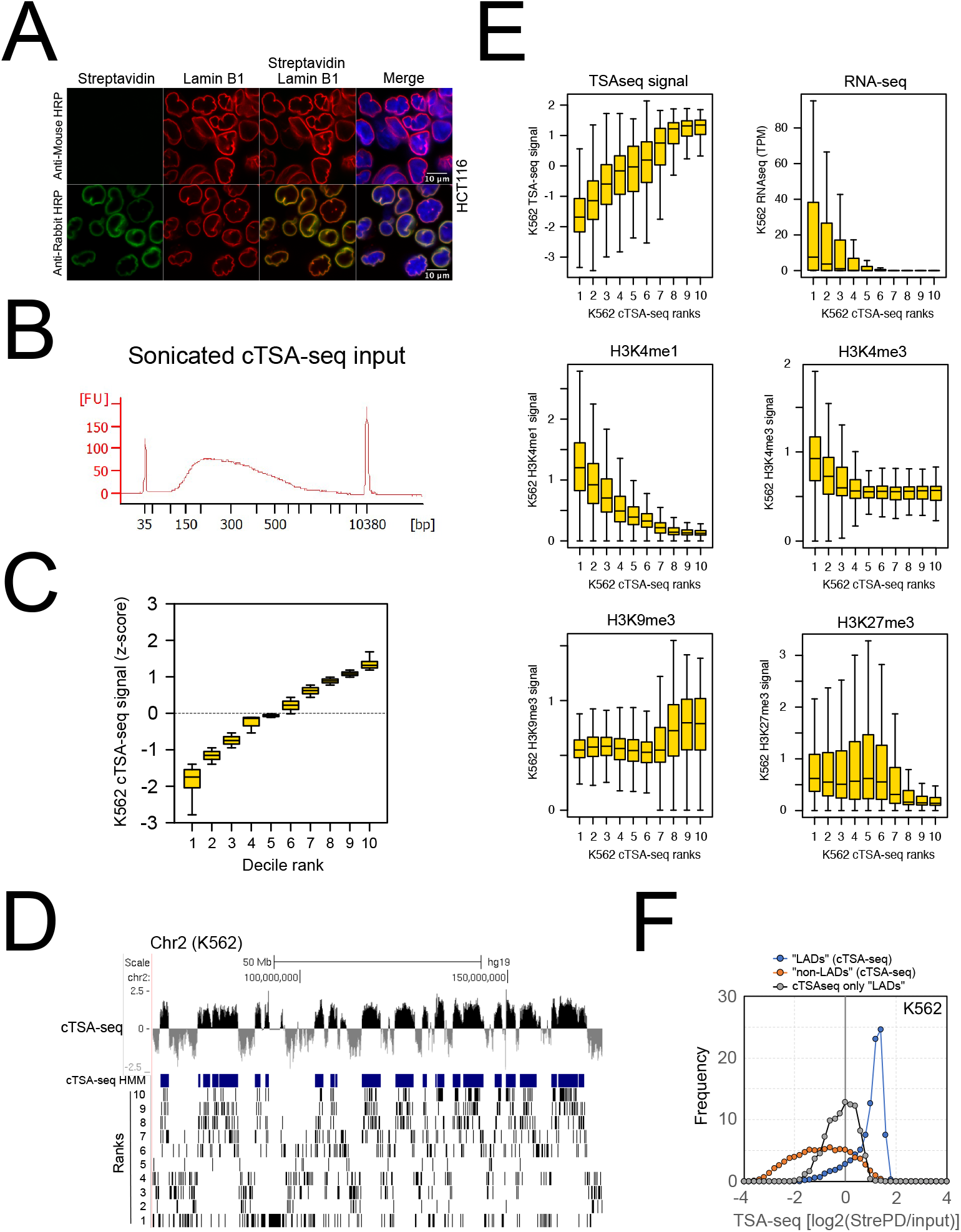
cTSA-seq labeling controls and additional details regarding cTSA-seq mapping. **(A)** The TSA reaction performed with HCT116 cells using an anti-rabbit HRP or an anti-mouse HRP secondary. Staining was done with Streptavidin (green) and lamin-B1 (red). **(B)** An example Bioanalyzer plot showing the distribution of DNA fragments in base pairs (“bp”) obtained after sonication. **(C)** Boxplot showing the distribution of K562 cTSA-seq signal in decile ranks. **(D)** UCSC Genome browser view (hg19) showing the location of decile ranks on chromosome 2. The decile ranks are show in descending order with the strongest cTSA-seq signal in rank 10 and the weakest in rank 1. The HMM track representing the called cTSA-seq NL-chromatin regions (“cTSA-seq HMM”) is shown in dark blue. **(E)** Boxplots showing the quantification of K562 TSA-seq (“TSA-seq signal”), RNA transcription (“RNA-seq”), euchromatin (“H3K4me1” and “H3K4me3”) and heterochromatin (“H3K9me3” and “H3K27me3”) signals across cTSA-seq deciles. **(F)** Lineplot showing the TSA-seq signal associated with different cTSA-seq features. We examined this signal over cTSA-seq defined regions that were colloquially called “LADs” (blue), “non-LADs” (orange) and “cTSA-seq only LADs” (grey).

**Figure S2:**
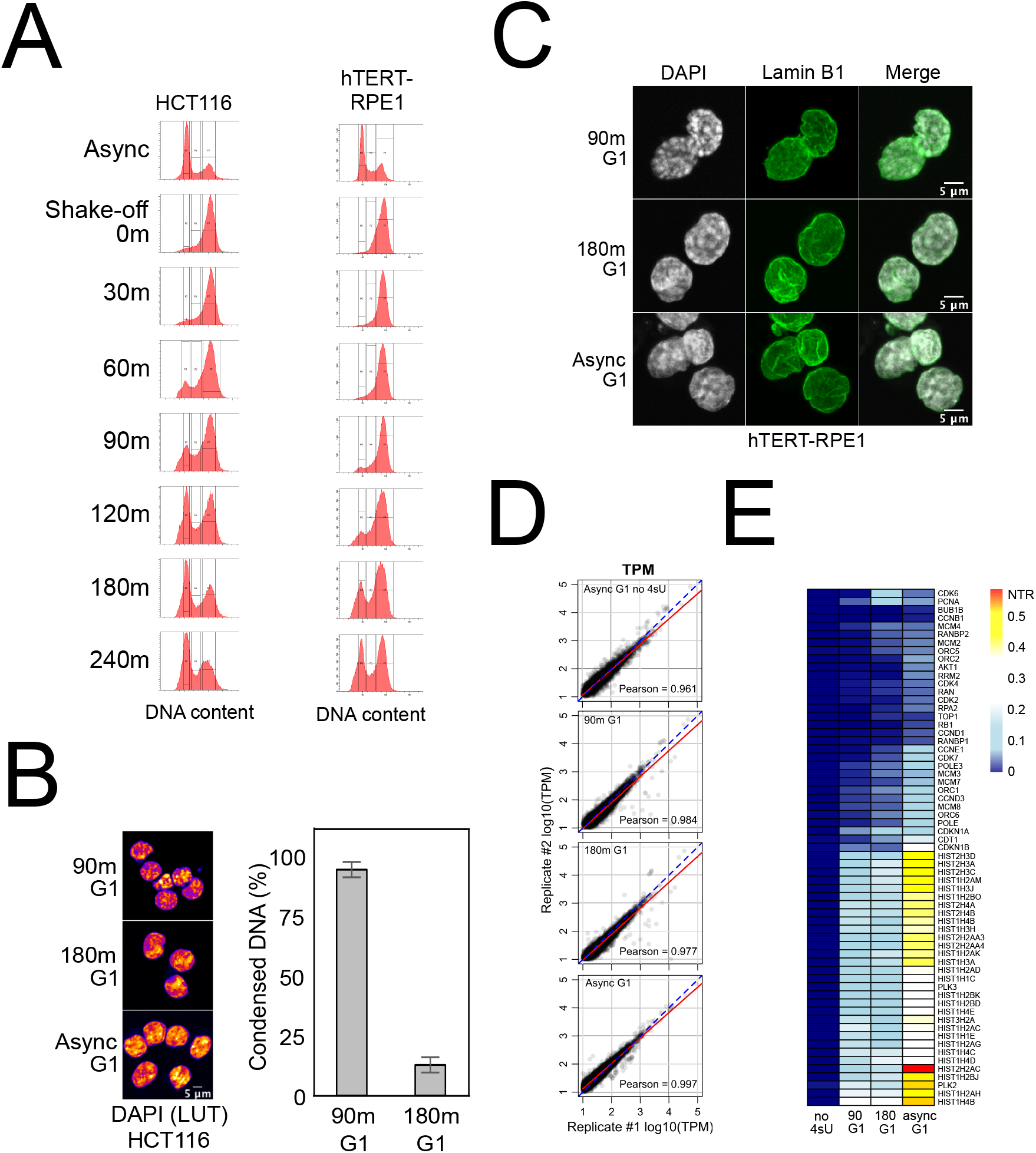
Quality control and features from early G1 FACS and SLAM-seq experiments. **(A)** Typical FACS profiles showing the time course of emergence of G1 cells after nocodazole release. DNA was stained with Hoechst 33342 prior to FACS. Asynchronous populations (“Async”) were used as a gating reference. The left and right panels are representative experiments done for HCT116 and hTERT-RPE1 cells, respectively. **(B)** Left panel shows a look up table (LUT) display of DAPI signal (DNA) for FACS sorted early G1 (90m G1), later G1 (180m G1) and asynchronous G1 (Async G1) HCT116 cells. Right panel shows a graph of the percentage of cells containing apparently dissimilar chromatin in the 90m early G1 population when compared to 180m G1 populations. **(C)** Lamin-B1 immunostaining for FACS isolated early (“90m G1”), later (“180m G1”) and asynchronous G1 (“Async G1”) hTERT-RPE1 cells. Lamin-B1 is presented in green and DNA (DAPI) is presented as grayscale. The scale bar represents 5 microns. **(D)** Scatterplots for SLAM-seq replicates. The axes are a log10 transformation of the transcripts per million (TPM). Untreated asynchronous G1 (“Async G1 no 4sU”), early (“90m G1”), later (“180m G1”) and asynchronous G1 (“Async G1”) HCT116 populations were examined. The blue dashed line is a reference diagonal line and the red line represents a linear model. The Pearson correlation value is presented in each panel on the lower right. **(E)** Heatmap showing the new transcription (NTR) level for select genes, including known cell cycle proteins and histone gene clusters, in the indicated cell populations. Note that the asynchronous G1 with no 4sU is labeled as “no 4sU”.

**Figure S3:**
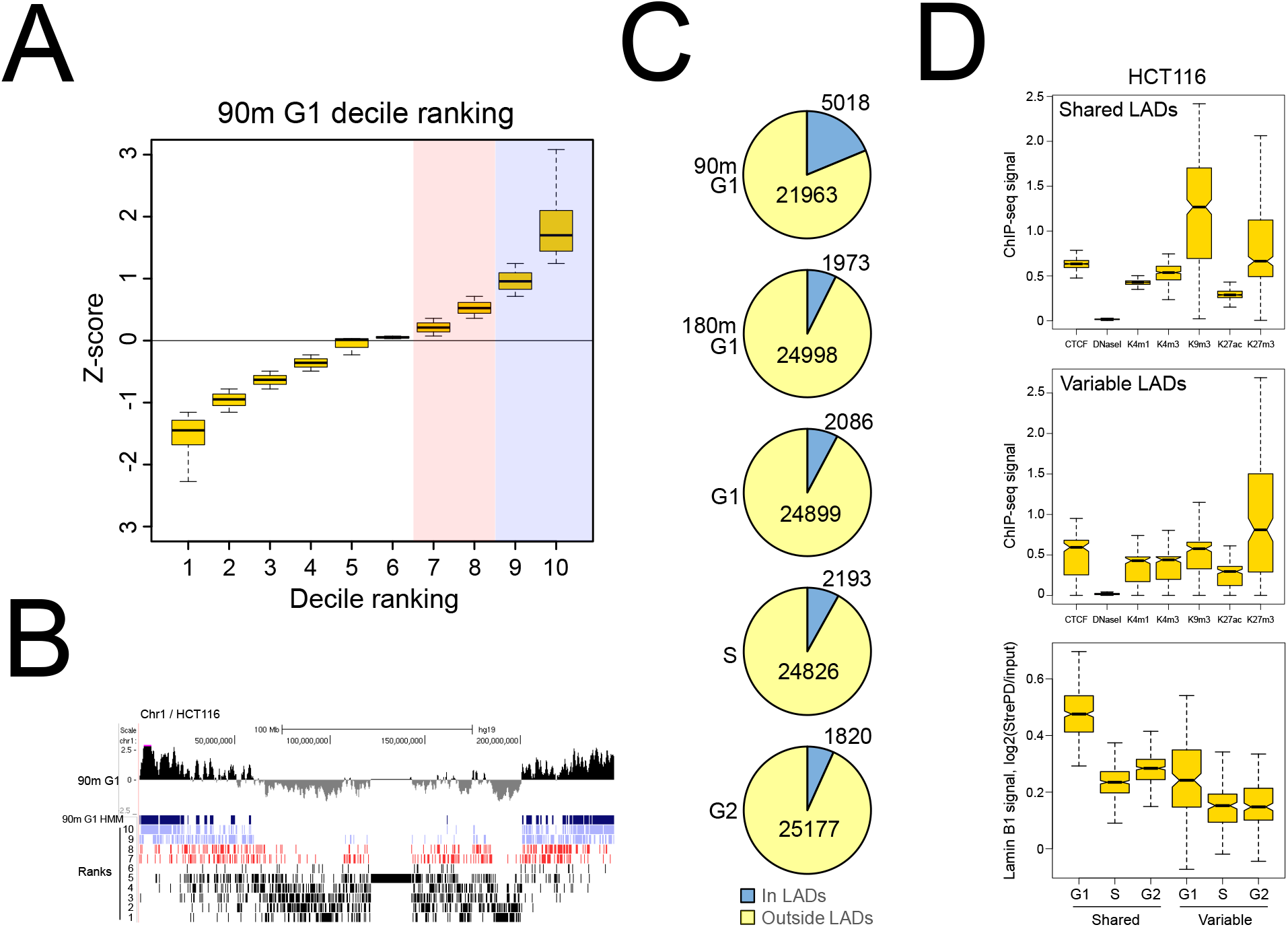
Decile ranking and additional analyses of cell cycle-related cTSA-seq experiments. **(A)** Example boxplot showing the decile ranking of cTSA-seq z-score data for the 90m G1 sample. The light red bar highlights a weak positive cTSA-seq z-score while the blue bar represents the strongest cTSA-seq z-score. **(B)** Example UCSC Genome Browser view (Chr1, hg19) showing the location of decile ranks for 90m G1 cells and their relationship to the HMM calls. The HMM call for NL-chromatin regions at 90m G1 is shown in dark blue. The 9^th^ and 10^th^ ranks, which correspond to the strongest cTSA-seq signal are shown in light blue and the 7^th^ and 8^th^ ranks are shown in red. **(C)** Pie-charts showing the number of genes inside and outside of the lamin-B1 cTSA-seq mapped regions (referred to here as “In LAD” and “Outside LAD”, respectively). Gene localization was called if at least 80% of the gene resided within the examined feature. **(D)** Boxplots showing the ENCODE HCT116 epigenetic ChIP-seq signal quantified over lamin-B1 cTSA-seq mapped chromatin regions that were shared (top panel, referred to as “Shared LADs”) or variable (middle panel, referred to as “Variable LADs”) across the G1, S, and G2 phases of the cell cycle. A boxplot showing the change in lamin-B1 cTSA-seq signal in shared and variable LADs over the cell cycle (bottom panel).

**Figure S4:**
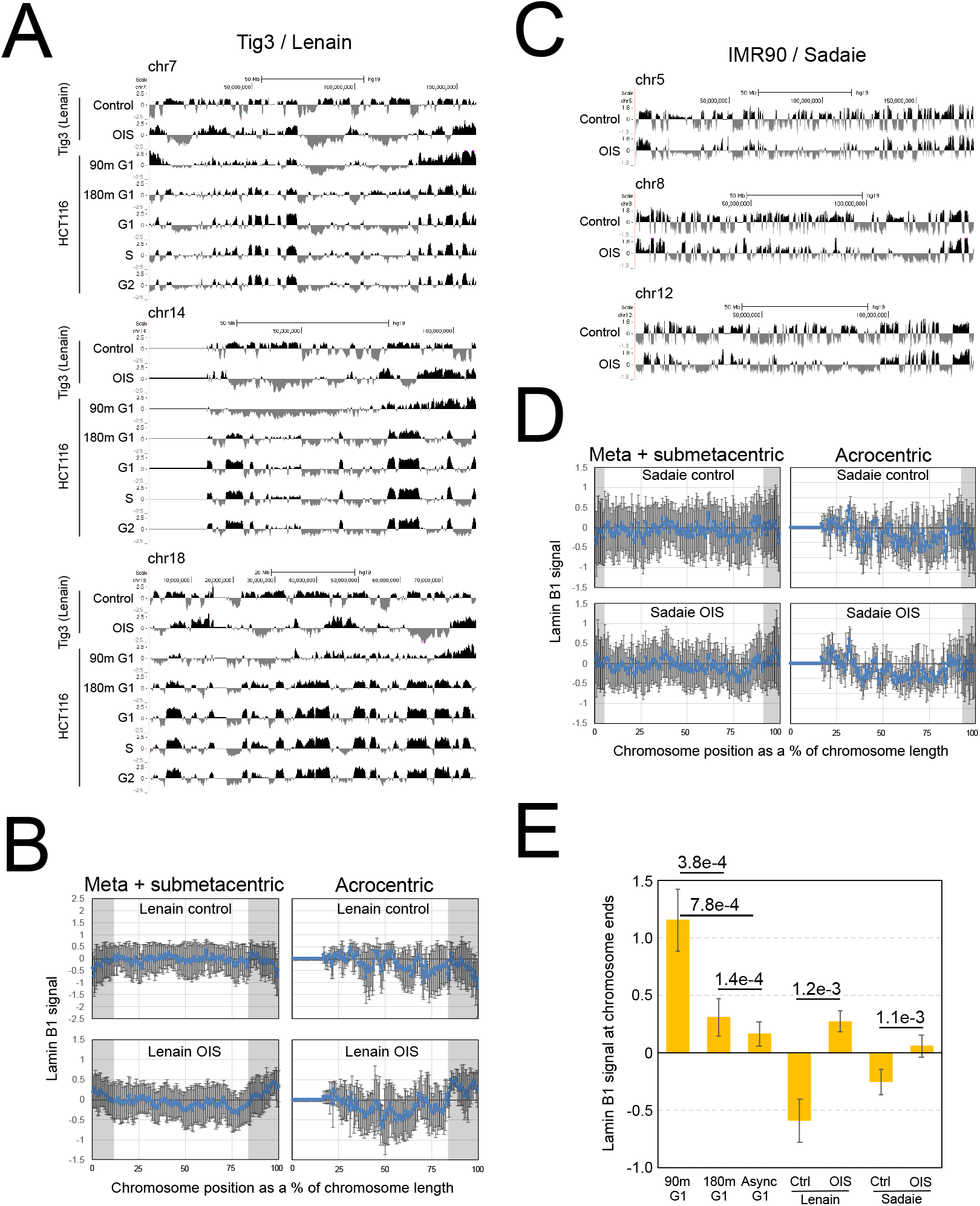
The early G1 chromatin at or near the reassembling NL identified by lamin-B1 cTSA-seq may be a feature present in Oncogene-Induced Senescence (OIS) cells. (**A**) UCSC Genome browser tracks (hg19) of LADs mapped by lamin-B1 DamID from “Control” and Oncogene-induced senescence (“OIS”) human Tig3 cells (29) and by lamin-B1 cTSA-seq from “90m G1”, “180m G1”, “G1”, “S” and “G2” HCT116 cells. Metacentric (chr 7), acrocentric (chr 14) and small metacentric (chr 18) chromosomes are shown. The cTSA-seq data presented are average z-scores. (**B**) Lineplots displaying the average lamin-B1 DamID signal from control and OIS Tig3 cells. Meta- and submetacentric (left) and acrocentric (right) chromosomes are shown. The error bars represent standard deviation. Grey bars highlight regions where the signal is elevated in OIS samples. (**C**) UCSC Genome browser tracks for lamin-B1 ChIP-seq signal (hg19) from “Control” and “OIS” IMR90 human cells (30). Chromosomes 5, 8 and 12 are presented. (**D**) Lineplots displaying the average lamin-B1 ChIP-seq signal from control and OIS IMR90 cells (30). Grey bars highlight regions where the signal is elevated in OIS cells. (**E**) Average HCT116 lamin-B1 cTSA-seq signal measured from the ends of chromosomes for 90m G1, 180m G1, asynchronous G1. The control (“Ctrl”) and OIS data are from previously published work shown in A-D (29,30). We defined the chromosome ends as the last 2.5% of each chromosome arm. The p-values represent are a two-tailed t-test and error bars represent the standard deviation of the signal.

